# Paired viromics resolves the modular ecological architecture of the swine nasopharyngeal phageome

**DOI:** 10.64898/2026.05.28.728360

**Authors:** Oscar Mencía-Ares, Carlus Deneke, Sonia Martínez-Martínez, Burkhard Malorny, César B. Gutiérrez-Martín, Josephine Gruetzke

## Abstract

**Background:** Bacteriophages are recognized modulators of microbiome composition and function, yet their role in the porcine upper respiratory tract, a primary gateway for pathogen colonization in the post-weaning period, remains unexplored. Unlike the porcine gut, no reference framework is available for respiratory sites. Furthermore, the low-biomass nature of nasopharyngeal specimens makes virome recovery highly sensitive to extraction strategy, but the extent to which workflow choice shapes ecological inference in this niche has not been evaluated.

**Results:** We profiled the nasopharyngeal phageome of post-weaning piglets across ten commercial farms (30 pen-level pools) using paired DNA-microbiome (DNA-m) and virus-like particle–enriched (VLP-e) short-read metagenomics (*n* = 60 libraries). Protocol choice strongly reshaped viral recovery (PERMANOVA R² = 0.448, *p* < 0.0001), with a contig overlap between workflows of <1%. DNA-m favored assembly contiguity, while VLP-e maximized viral detection. By integrating both approaches, we constructed a curated catalogue of 2,501 non-redundant viral operational taxonomic units (vOTUs), with only 5.2% showing similarity to known phages, underscoring the extensive novelty of this niche. Ecologically, within the integrated community dataset (*n* = 4,357), predicted replication strategy emerged as a dominant organizing axis: lifestyle explained up to 40.6% of compositional variation at family level. Host prediction linked phages to dominant upper-airway colonizers, including *Streptococcaceae*, *Moraxellaceae*, *Pasteurellaceae*, with a marked lifestyle–host polarization: virulent phages were preferentially linked to *Bacteroidota* (particularly *Prevotella*), whereas temperate phages were enriched in *Streptococcaceae* and *Moraxellaceae*. Integration of viral taxonomy and host affiliation resolved a modular architecture in which a few recurrent phage–host couplings (e.g., *Suoliviridae–Bacteroidota*, *Peduoviridae–Pasteurellaceae*, *Aliceevansviridae–Streptococcaceae*) were conserved but differentially weighted between virulent and temperate fractions.

**Conclusions:** This study establishes the first phageome catalogue and ecological framework for a respiratory site in livestock. The nasopharyngeal phageome is organized into recurrent, host-linked taxonomic modules jointly constrained by viral lineage, host affiliation and replication strategy, with lifestyle-dependent connections to key colonizers implicated in the porcine respiratory disease complex. This catalogue and its modular architecture provide a foundation for investigating phage-mediated modulation of bacterial dynamics during the post-weaning transition and for the selection of lytic phage candidates targeting respiratory pathogens.

## Background

The porcine respiratory disease complex (PRDC) is among the most important health challenges in intensive pig production, particularly during the post-weaning period [1,2]. Although the clinical and economic dimensions of PRDC are well documented [3], the microbial ecology of the porcine upper respiratory tract (a primary gateway for pathogen colonization and transmission) remains incompletely understood [4]. To date, culture-independent profiling of this niche has focused almost exclusively on bacteria [5,6], leaving the viral component largely unexplored. This gap is notable because bacteriophages (viruses that infect bacteria) are now recognized as ubiquitous modulators of microbiome composition and function across host-associated ecosystems [7,8].

Phages shape bacterial communities through two contrasting replication strategies: virulent phages kill their hosts and can promote bacterial turnover, whereas temperate phages integrate as prophages and reshape host fitness through lysogenic conversion and horizontal gene transfer [9,10]. Because these lifestyles generate distinct phage–host association patterns [11,12], replication strategy has emerged as a key ecological axis in phageome research.

Viral metagenomics has dramatically expanded the known phage diversity in animal-associated microbiomes [13,14]. In pigs, however, phageome characterization has concentrated on the gastrointestinal tract, where high microbial and viral biomass and relatively mature viromics workflows have enabled the construction of large-scale reference databases [15,16]. By contrast, the porcine respiratory phageome has not been systematically characterized.

This gap is compounded by the technical constraints of respiratory viromics: upper respiratory tract specimens are low-biomass samples in which host DNA dominates sequencing libraries [5], severely diluting viral signal. Two main strategies exist for recovering viral sequences from such samples: DNA microbiome shotgun metagenomics (DNA-m), which sequences all extracted nucleic acids, and virus-like particle enrichment (VLP-e), which uses filtration and nuclease treatment to deplete cellular DNA prior to sequencing. These approaches differ in signal-to-noise ratio and can recover partially non-overlapping viral fractions [17,18]; however, their side-by-side performance in swine respiratory specimens has not been evaluated.

Here, we address these gaps using short-read metagenomics to profile the nasopharyngeal phageome of post-weaning piglets from ten Spanish commercial farms. Through a paired design in which the same 30 pen-level samples were processed with both DNA-m and VLP-e workflows (*n* = 60 libraries), we quantify how protocol choice influences viral recovery and community structure. By integrating both workflows, we construct a curated, non-redundant catalogue of respiratory viral operational taxonomic units (vOTUs) and examine how predicted replication strategy and host affiliation organize the nasopharyngeal phageome, supported by complementary phylogenetic and CRISPR-based host-linking evidence.

## Results

### Protocol-dependent recovery of complementary nasopharyngeal phageome fractions

Paired nasopharyngeal swabs from post-weaning piglets across ten intensive farms (30 sampling points; *n* = 60 libraries) were processed using either a DNA microbiome protocol (DNA-m) or a virus-like particle–enriched protocol (VLP-e) to quantify how extraction strategy influences nasopharyngeal phageome recovery. The complete experimental and bioinformatic workflow is summarized in Figure 1.

**Figure 1.**
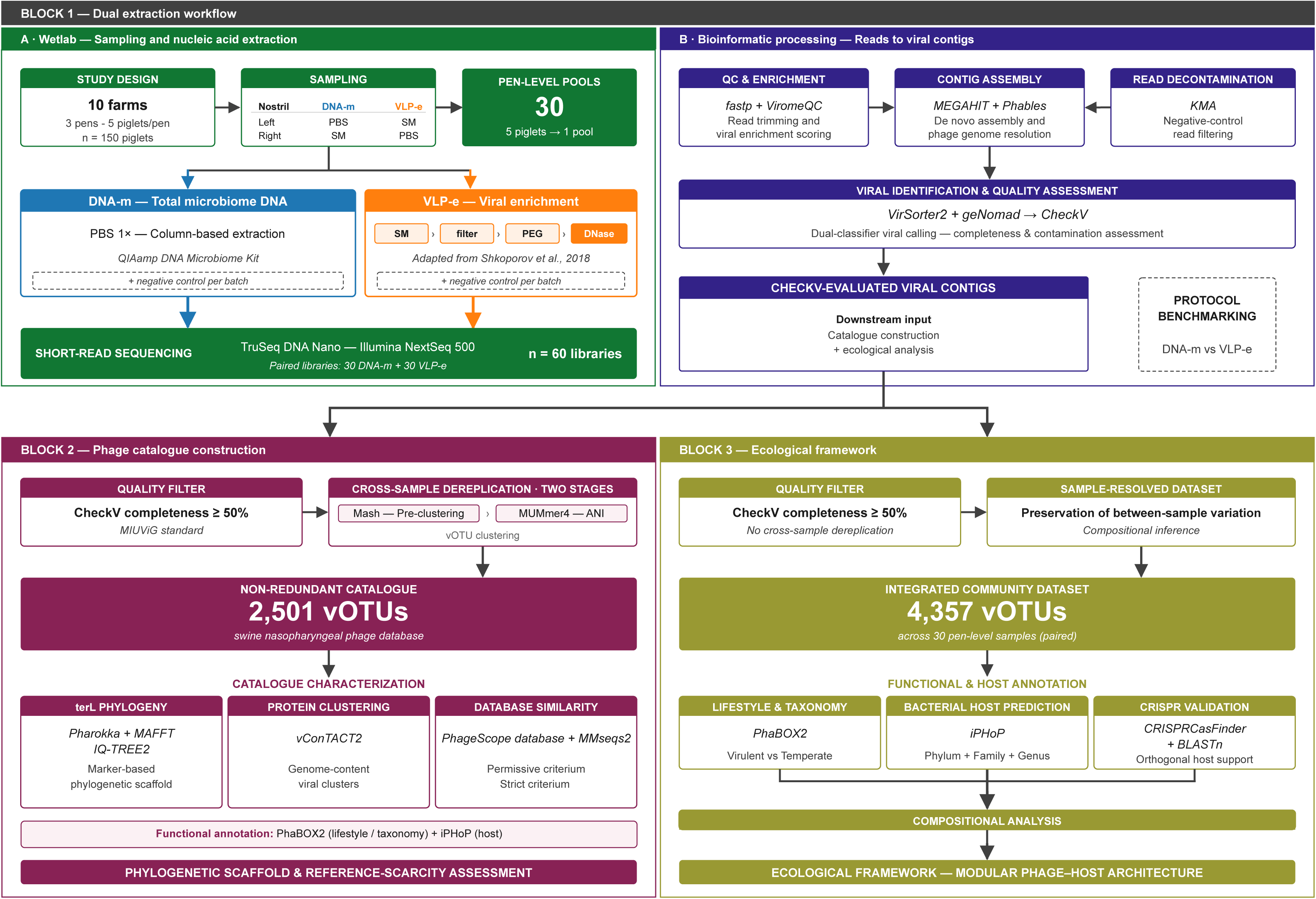
Overview of the experimental and bioinformatic workflow. Paired nasopharyngeal swab pools from post-weaning piglets (*n* = 30 pen-level samples from ten commercial farms) were processed through two complementary extraction protocols (DNA-m and VLP-e; *n* = 60 libraries), integrated into a curated catalogue of 2,501 representative vOTUs, and analyzed within an ecological framework combining predicted lifestyle, viral taxonomy, and bacterial host affiliation.

Upstream sequencing and enrichment metrics indicated that the two approaches favor distinct fractions of the sample. DNA-m libraries yielded more reads per sample than VLP-e (50.2 ± 9.0 vs 32.0 ± 17.1 million; *p* < 0.0001; Figure 2A). In contrast, VLP-e showed markedly improved viral enrichment, with higher ViromeQC enrichment scores (*p* < 0.0001; Figure 2B) and reduced relative alignment to rRNA and bacterial marker genes (all *p* < 0.001; Additional file 1 – Table S1), supporting effective depletion of cellular DNA. These differences were mirrored in assembly profiles, with VLP-e producing more contigs and DNA-m producing longer contigs on average (both *p* < 0.0001; Figure 2C-D; Additional file 1 – Tables S2-S3).

**Figure 2.**
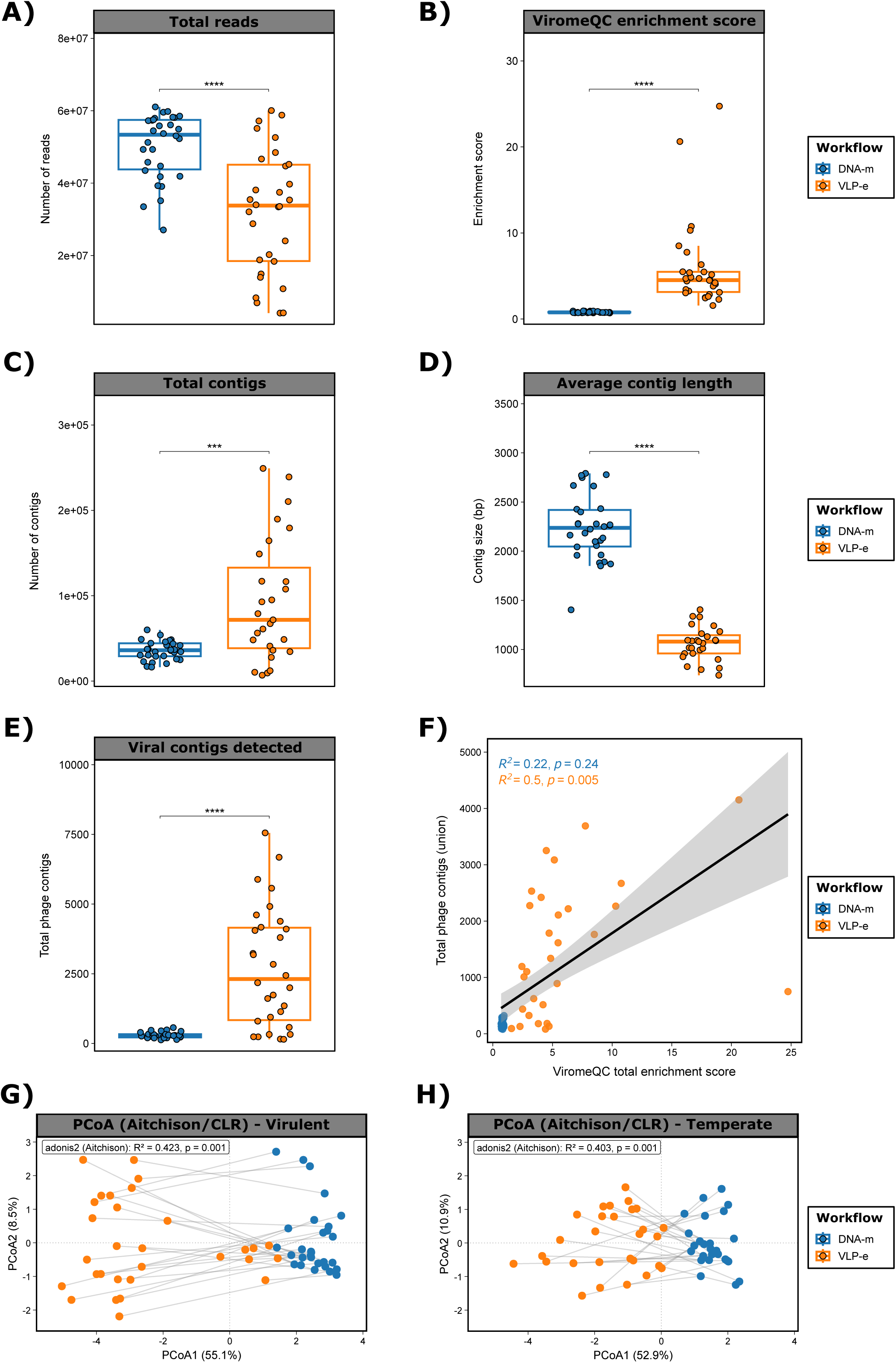
Protocol-dependent recovery of paired nasopharyngeal phageome fractions. **A)** Total sequencing reads (bp) per sample. **B)** ViromeQC total enrichment score. **C)** Total number of assembled contigs. **D)** Average contig length (bp). **E)** Total phage contigs detected. In panels A–E, each sample is represented by a dot with horizontal jitter for visibility. Horizontal box lines represent the first quartile, the median and the third quartile; whiskers extend to 1.5× the interquartile range. Differences between paired samples were evaluated with the Wilcoxon signed-rank test. Asterisks denote statistical significance: *** *p* < 0.001; **** *p* < 0.0001. **F)** Spearman correlation between ViromeQC total enrichment score and the number of phage contigs recovered per sample, stratified by extraction method. Regression lines and 95% confidence intervals are shown. **G)** Principal coordinates analysis (PCoA) based on Aitchison distances (CLR-transformed vOTU abundances) at family level for the virulent and H) **temperate** phage fraction, colored by extraction method. PERMANOVA R² and *p*-value are indicated. DNA-m: DNA microbiome protocol; VLP-e: virus-like particle–enriched protocol. *n* = 60 libraries (30 DNA-m + 30 VLP-e).

To evaluate whether enrichment translated into improved viral recovery, assembled contigs were screened with VirSorter2 and geNomad. Across classifiers, VLP-e assemblies consistently yielded more viral contigs than DNA-m assemblies (*p* < 0.0001; Figure 2E), and this advantage remained after normalization for assembly size and contig number (*p* < 0.001), indicating that improved detection was not driven solely by assembly output. ViromeQC enrichment scores were positively associated with the number of viral contigs detected (R^2^ = 0.5, *p* < 0.01; Figure 2F), further linking enrichment performance to viral recovery.

Finally, protocol choice reshaped the recovered viral community rather than producing a simple quantitative shift. When viral contigs were dereplicated into viral operational taxonomic units (vOTUs), overlap between paired analyses was minimal, with >99% of vOTUs detected exclusively by one protocol. Consistently, CLR/Aitchison-based analyses showed a strong extraction-method effect on community structure (PERMANOVA, R² = 0.448, *p* < 0.0001), which persisted when stratified by lifestyle (virulent R² = 0.423; temperate R² = 0.403; both *p* < 0.0001) and was reflected by clear protocol-driven separation in PCoA ordinations (Figures 2G-H). Together, these results indicate that VLP-e and DNA-m recover largely non-overlapping components of the nasopharyngeal phageome, supporting their complementary use to obtain a more comprehensive view of viral diversity in the porcine upper respiratory tract.

### Construction of a curated non-redundant swine nasopharyngeal phage database

To enable comparative analyses of swine nasopharyngeal viromes, we built a curated, non-redundant database integrating viral sequences recovered across both extraction protocols. After merging geNomad and VirSorter2 predictions and resolving inter-tool redundancies, the initial integrated set comprised 51,175 viral contigs, with substantial agreement between classifiers (Additional file 2 – Figure S1A). CheckV quality assessment categorized most predicted viral contigs as low-quality (83.7%), with medium-quality (5.3%), high-quality (3.6%), and complete genomes (2.0%) representing smaller fractions. Viral contigs of medium quality or higher (*n* = 8,854) were retained and dereplicated by sequence similarity to yield a final non-redundant catalogue of 2,501 representative vOTUs, hereafter referred to as the swine nasopharyngeal phage database released with this study (https://doi.org/10.5281/zenodo.20119467).

The resulting catalogue captured largely novel viral diversity. Only 130 representative vOTUs (5.2%) met the permissive criteria for similarity to reference phage sequences (for details see Methods), and only 33 (1.3%) of these met the stringent high-confidence threshold. This direct sequence-similarity assessment indicates that swine nasopharyngeal phages are scarcely represented in available reference databases.

At a high level, the curated database was dominated by tailed dsDNA bacteriophages, with 98.9% of representative vOTUs assigned to both the *Uroviricota* phylum and the *Caudoviricetes* class. Taxonomic resolution decreased substantially at lower ranks, barely reaching family-level assignment for only 10.1% of vOTUs. Among assigned viral families, *Salasmaviridae* emerged as the most frequently detected (21.0%), followed by *Autographiviridae* (11.5%), *Rountreeviridae* (11.5%), and *Steigviridae* (10.7%). Lifestyle prediction indicated a predominance of virulent phages (*n* = 1,335; 53.4%), followed by temperate phages (*n* = 928; 37.1%), with 9.5% remaining unclassified. Interestingly, predicted proviruses were infrequent (2.8%). Bacterial host prediction remained unresolved for approximately half of the representative vOTUs (*n* = 1,283; 51.3%); among assigned hosts, *Bacillota* predominated (29.3%), followed by *Bacteroidota* (7.6%) and *Pseudomonadota* (6.7%).

To resolve lineage structure independently of database-anchored taxonomy, we used the large terminase subunit (terL) as a conserved phylogenetic marker. A total of 1,449 representative vOTUs (57.9%) encoded a terL of sufficient length and were placed on a maximum-likelihood phylogeny (Figure 3). Hierarchical clustering of cophenetic distances on this tree partitioned terL-positive vOTUs into 19 lineages, with markedly uneven sizes, with the four largest clusters accounting for >70% of all terL-encoding vOTUs, while the remaining clusters contained smaller or rare lineages. Annotation of each cluster with viral taxonomy, predicted lifestyle, and predicted host range showed coherent assignment patterns, with vOTUs from the same viral family generally co-clustering closely (Additional file 3).

**Figure 3.**
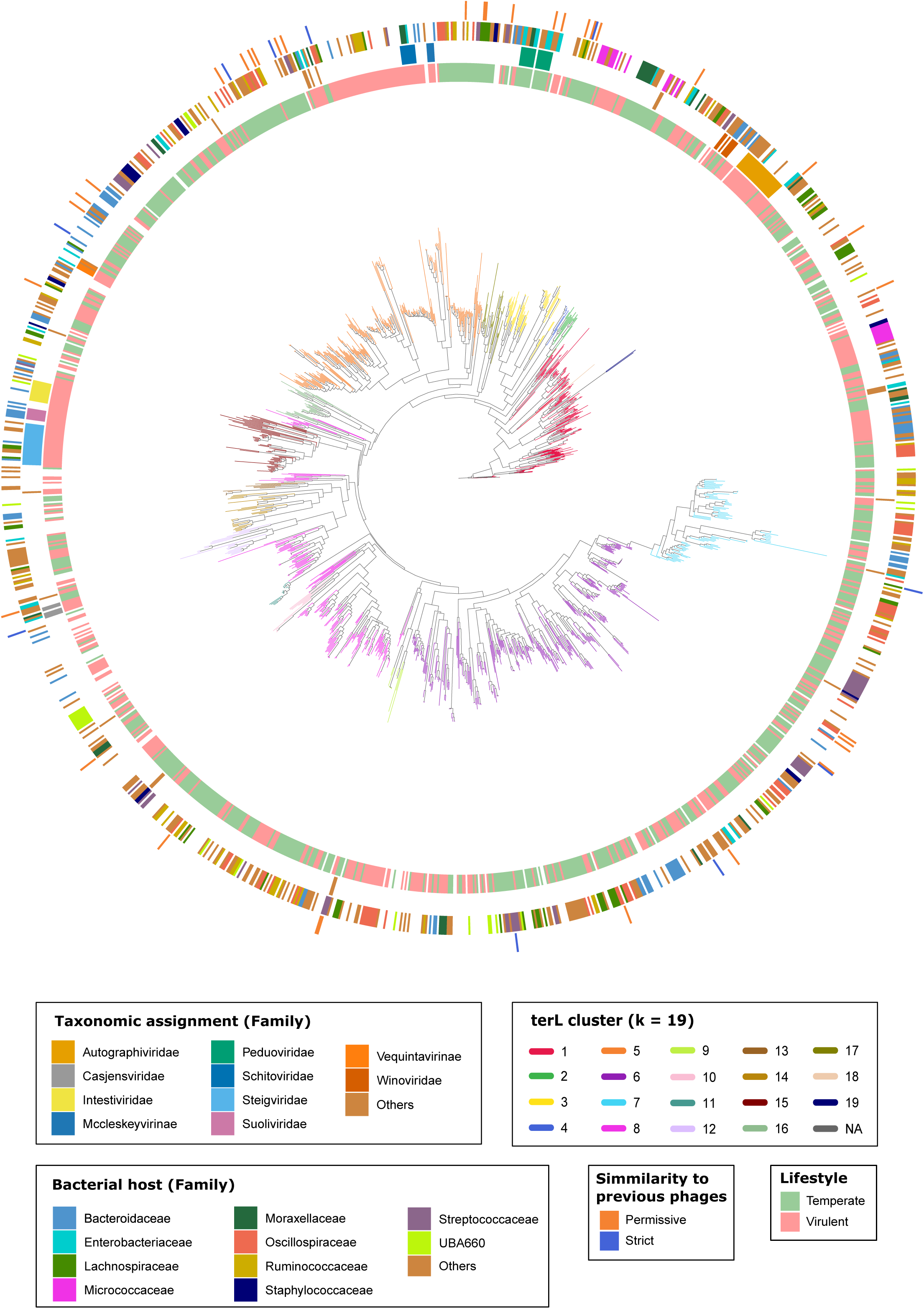
Phylogenetic coherence of the curated vOTU catalogue based on the large terminase subunit (terL). Maximum-likelihood circular phylogeny of terL amino-acid sequences (≥100 residues) extracted from representative vOTUs, inferred with IQ-TREE2 under the LG+G4 model. *n* = 1,449 representative vOTUs grouped into 19 clusters by hierarchical clustering of cophenetic distances. Concentric annotation rings display, from inner to outer: predicted lifestyle, predicted host family, similarity to previously described phage sequences, and viral family assignment. Cluster identifiers are labelled at the corresponding tree sections.

To complement the single-marker terL phylogeny with a whole-genome, protein-content-based view of phage relatedness, we processed the catalogue jointly with reference genomes through vConTACT2, which partitions phage genomes into viral clusters (VCs). The catalogue was distributed across 392 VCs of ≥2 vOTUs, ranging from 2 to 94 vOTUs in size. Within the network, 1,021 vOTUs (40.8%) co-clustered with at least one reference genome and shared protein-content architecture with previously described viruses, although only 279 (11.2% of the catalogue) co-clustered with a reference genome carrying a formal taxonomic assignment. The remaining 1,480 vOTUs (59.2%) had no detectable analogue in current reference collections: 645 (25.8%) clustered exclusively with other catalogue vOTUs, including 146 VCs of related sequences internal to the nasopharyngeal phageome, while 835 (33.4%) remained as singletons. A detailed description of these VCs is available in Additional file 4 (Table S7).

Direct comparison of the two clustering schemes confirmed their complementary, hierarchically nested nature. The asymmetric adjusted Wallace coefficients revealed a strongly nested structure: VC membership was highly predictive of terL co-clustering (W_VC→terL_ = 0.86; 95% CI 0.84–0.89), whereas the converse was not (W_terL→VC_ = 0.04; 95% CI 0.03–0.05), indicating that vConTACT2 clusters resolve into finer relationships embedded within broader terL lineages. Consistently, the global adjusted Rand index between the two partitions was low (ARI = 0.074), as expected when one classification subdivides another. Interestingly, of the 2,501 vOTUs, 1,018 were placed in a cluster by both methods, 730 only by vConTACT2, 418 only by terL, and 335 by neither (Additional file 4 – Tables S8-S9).

To describe the ecological footprint of the curated catalogue across the sampled population, we extended the analysis from the 2,501 representative vOTUs to all detected member sequences within their respective vOTU clusters across the 30 pen-level samples (Additional file 2 – Figure S1B). Cross-sample prevalence, defined as the number of samples in which a particular vOTU was detected, followed a strongly long-tailed distribution: 1,820 vOTUs (72.8%) were detected in a single sample, 656 (26.2%) in two or more, 101 (4.0%) in five or more, and only 26 (1.0%) in ten or more samples; no vOTU was detected in all 30 samples. Within-sample multiplicity, defined as the count of a given vOTU within a sample across both extraction protocols, was equal to one (e.g. single occurrence of a given vOTU in a sample) for the majority of vOTU–sample observations across both extraction protocols, with multiplicities ≥ 2 occurring infrequently and concentrated among vOTUs of higher cross-sample prevalence.

Finally, to assess whether the present sampling effort had captured the recoverable catalogue diversity, we built a sample-based vOTU accumulation curve from random permutations of sample order. Here, we observed that the curve continued to rise at *n* = 30 without inflection toward an asymptote (Additional file 2 – Figure S1C). A power-law fit was decisively preferred over an asymptotic exponential by AIC (ΔAIC = 391), confirming that vOTU richness has not saturated at the current sampling depth and is consistent with an open viral pool. Together, these patterns indicate that the 2,501 representative vOTUs constitute a partial yet informative view of the swine nasopharyngeal phageome, providing a sufficient basis to characterize its broad compositional and functional features at the population level.

### Global characterization of the swine nasopharyngeal phageome

#### Lifestyle is a major organizing axis of the nasopharyngeal phageome

To characterize sample-level organization of the swine nasopharyngeal phageome, we analyzed an integrated community dataset comprising all vOTUs with CheckV-estimated completeness ≥ 50% detected across both extraction protocols (*n* = 4,357), thereby preserving within-sample diversity and between-sample compositional variation for ecological inference. We then tested whether phage lifestyle acts as a primary organizing axis and whether this structure resolves into recurrent host–taxonomy modules.

At a broad taxonomic level, the community was dominated by tailed dsDNA phages, with *Caudoviricetes* class accounting for 99.2% of vOTUs. Taxonomic resolution decreased sharply at lower ranks, with 90.2% and 80.8% of vOTUs remaining unclassified at the family and genus levels, respectively. Within the classified fraction, a small set of families concentrated most assigned abundance, including *Salasmaviridae*, *Peduoviridae*, *Rountreeviridae* and *Autographiviridae* (Figure 4A), indicating a highly diverse but sparsely classified community. A detailed summary of viral taxonomy at family and genus level is available in Additional files 5 and 6 (Figure S2A).

**Figure 4.**
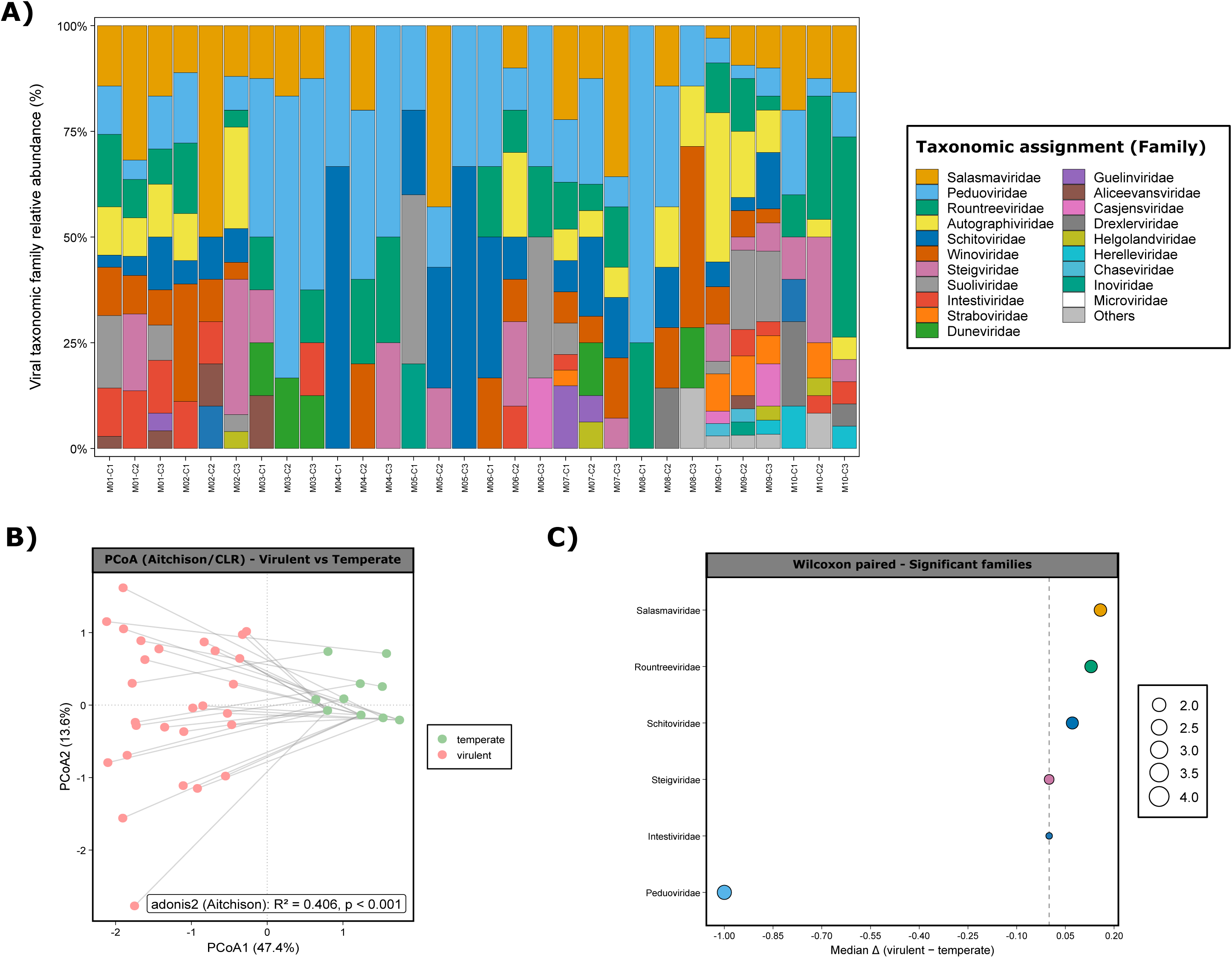
Lifestyle as a major organizing axis of the nasopharyngeal phageome. **A)** Stacked bar plot of the proportion (%) of vOTUs counts classified by viral family per sample, ordered by farm and pen. Colors represent individual viral families. **B)** Principal coordinates analysis (PCoA) based on Aitchison distances (CLR-transformed vOTU abundances) aggregated at family level, colored by predicted lifestyle. PERMANOVA R² and *p*-value are shown. **C)** Dot plot of viral families with significant lifestyle-associated differential abundance (*p* < 0.05; paired Wilcoxon signed-rank test). Points indicate the median paired difference (Δ) between virulent and temperate CLR-transformed abundances; negative values indicate temperate enrichment, and positive values indicate virulent enrichment. Point size reflects the –log10 adjusted *p*-value. Analyses were performed on the integrated community dataset (*n* = 4,357 vOTUs with CheckV-estimated completeness ≥50%) across 30 pen-level samples.

Although taxonomic resolution was limited at lower ranks, lifestyle-based comparisons still revealed a strong and consistent structuring signal. CLR/Aitchison-based PERMANOVA revealed strong lifestyle-associated differences at both the family (R² = 0.406; *p* < 0.0001; Figure 4B) and genus levels (R² = 0.612; *p* < 0.0001). Differential analyses identified 6 families with significant lifestyle biases (Figure 4C; Additional file 5 – Table S12), with *Peduoviridae* markedly enriched in temperate fractions and *Salasmaviridae*, *Rountreeviridae* and *Schitoviridae* enriched in virulent fractions. Differential results at genus level are available in Additional file 5 (Tables S13-S15).

Together, these results indicate that lifestyle rebalances the representation of specific viral lineages, generating distinct virulent- and temperate-associated community profiles.

#### Host affiliations are structured by lifestyle and polarized across dominant respiratory taxa

Bacterial host prediction remained unresolved for 50.3% of vOTUs. Among assigned hosts, phages were predominantly linked to *Bacillota* (33.1%), *Bacteroidota* (19.9%) and *Pseudomonadota* (19.3%). Predicted hosts included lineages typical of the upper respiratory tract, with *Streptococcaceae*, *Moraxellaceae* and *Pasteurellaceae* among the most represented families (Figure 5A), and *Streptococcus* (9.1%), *Prevotella* (8.2%), *Moraxella* (5.9%) and *Glaesserella* (2.6%) among the most frequent genera (Additional file 6 – Figure S2B).

**Figure 5.**
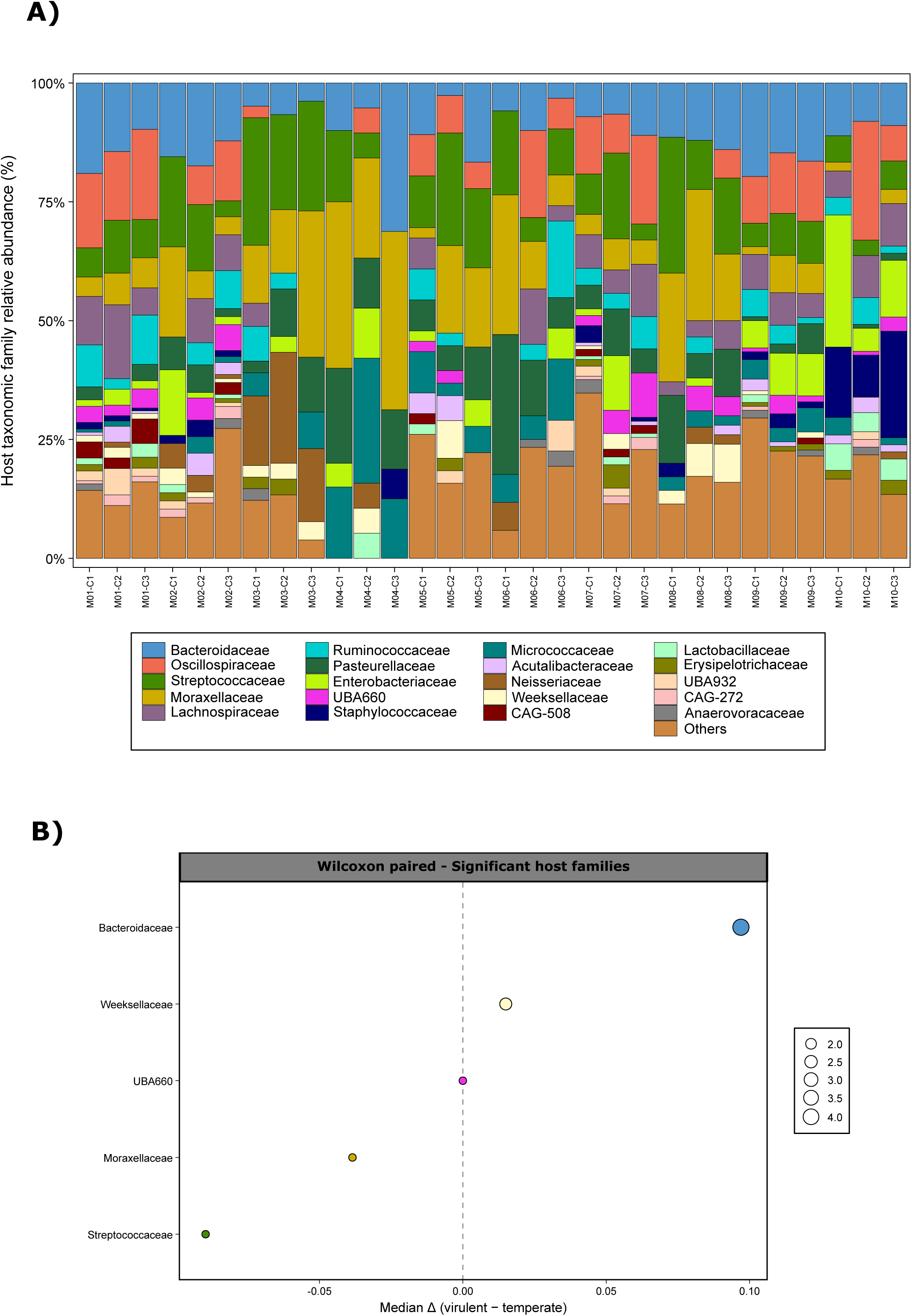
Host affiliations are structured by lifestyle and polarized across dominant respiratory taxa. **A)** Stacked bar plot of the proportion (%) of predicted host families (iPHoP) of vOTUs counts per sample, excluding vOTUs without host assignment. Samples are ordered by farm and pen. **B)** Dot plot of host families with significant lifestyle-associated differential abundance (*p* < 0.05; paired Wilcoxon signed-rank test on CLR-transformed abundances). Points indicate the median paired difference (Δ) between virulent and temperate fractions; positive values indicate virulent enrichment, and negative values indicate temperate enrichment. Analyses were performed on the integrated community dataset (*n* = 4,357 vOTUs) across 30 pen-level samples.

Host assignment rates differed markedly by lifestyle: temperate phages were substantially more likely to receive a host prediction than virulent phages (68.9% vs 38.9%; OR = 3.49; *p* < 0.0001). Beyond assignment probability, lifestyle also structured host composition. PERMANOVA detected moderate lifestyle-associated shifts in predicted host profiles at both the phylum (R² = 0.134; *p* < 0.0001) and family levels (R² = 0.072; *p* < 0.0001), indicating that lifestyle-dependent host partitioning is detectable across taxonomic scales.

Directional analyses clarified the polarity of this partitioning (Figure 5B). Virulent phages were preferentially associated with *Bacteroidota* (median Δ = 0.162; *p* < 0.001), particularly *Bacteroidaceae*, largely driven by *Prevotella*. In contrast, temperate phages were preferentially associated with *Bacillota* (median Δ = −0.137; *p* < 0.001) and *Pseudomonadota* (median Δ = −0.089; *p* < 0.001), mainly through enrichment in *Streptococcaceae* and *Moraxellaceae* and their corresponding genera *Streptococcus* and *Moraxella*. Detailed information is available at the different taxonomic levels in Additional file 7.

#### CRISPR links independently support host assignments without lifestyle bias

To independently support predicted host calls and the lifestyle-dependent host structuring described above, we examined CRISPR spacer–protospacer matches (Additional file 8). We identified 102 spacer-like matches corresponding to 65 vOTUs. For the 58 vOTUs supported by both CRISPR and iPHoP, host assignments were highly concordant (98.3% at phylum and 90.6% at family level), with substantial agreement at the genus level (67.9%). CRISPR-supported hosts were dominated by upper respiratory tract-associated taxa, particularly *Micrococcaceae* (driven by *Rothia*), with additional support for *Moraxellaceae* and *Pasteurellaceae*. The lifestyle distribution of CRISPR-supported phages did not differ from that of the full dataset.

#### Host–taxonomy coupling resolves into recurrent ecological modules

Finally, we integrated viral taxonomy, host range and lifestyle using enrichment-based host–phage association analyses. The nasopharyngeal phageome displayed a highly structured and non-random organization, resolving into a limited number of recurrent host-linked modules that were conserved across lifestyles but differentially emphasized between virulent and temperate fractions.

At the phylum level, three dominant axes captured most significant associations (*p* < 0.0001): *Suoliviridae* were strongly linked to *Bacteroidota* (OR = 255.8), *Peduoviridae* to *Pseudomonadota* (OR = 334.3), and *Rountreeviridae* to *Bacillota* (OR = 66.3) (Additional file 9). These significant modules persisted after lifestyle stratification and became more pronounced at finer host resolution (Figure 5; Additional files 10-11). Remarkably, within the virulent fraction, the *Suoliviridae*–*Bacteroidota* axis was largely driven by *Bacteroidaceae*, particularly *Prevotella*, while additional virulent-associated modules included enrichment of *Schitoviridae* in *Moraxellaceae* and *Pasteurellaceae* (Figure 6A-B). Conversely, temperate phages showed strong and specific links within respiratory-associated lineages, including enrichment of *Peduoviridae* in *Pasteurellaceae*, and highly specific associations between *Aliceevansviridae* and *Streptococcaceae* (Figure 6C-D). Extended enrichment statistic summaries are provided at phylum, family and genus level in Additional files 9-11.

**Figure 6.**
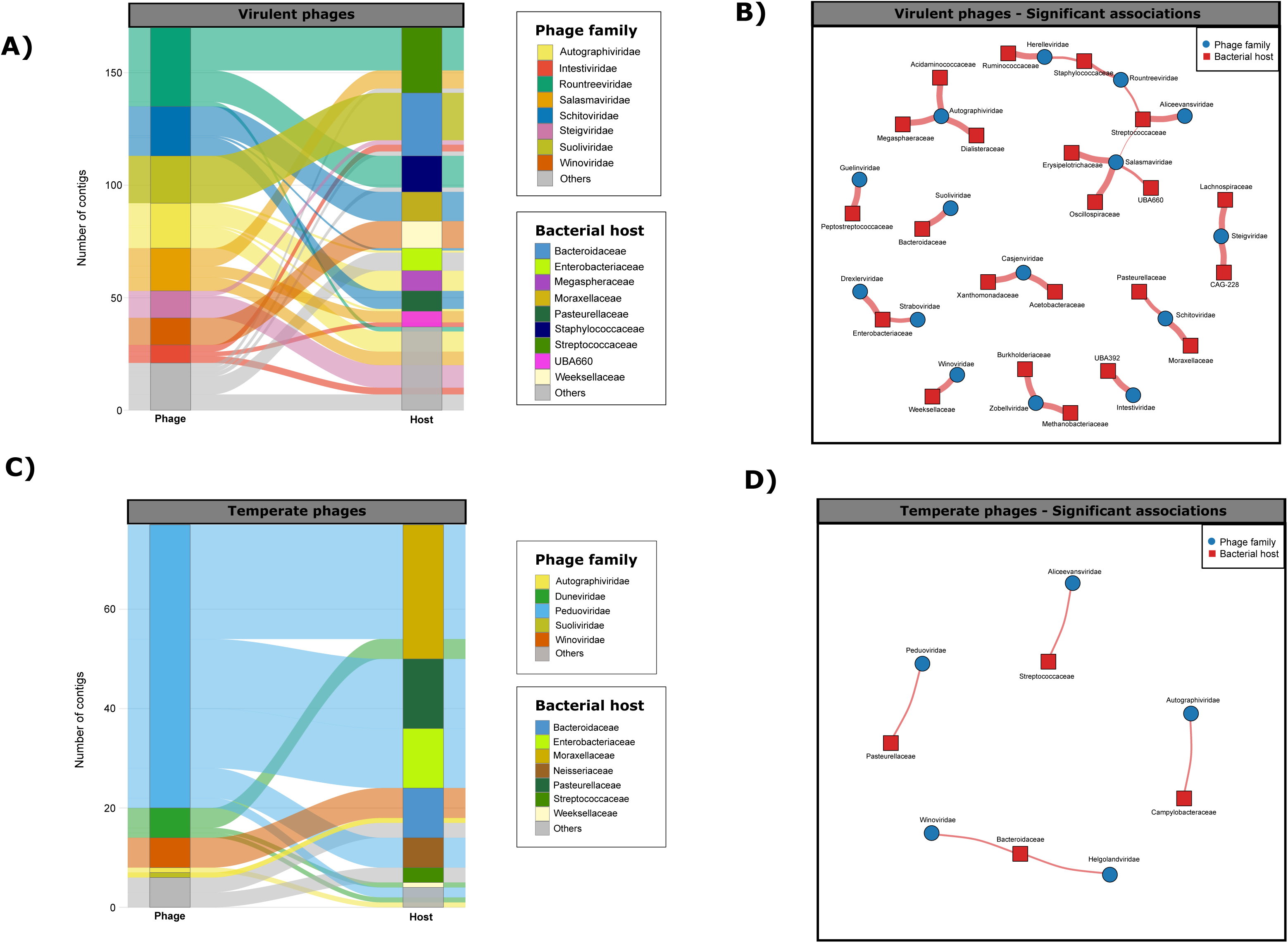
Host–taxonomy coupling resolves into recurrent ecological modules stratified by lifestyle. **A)** Alluvial plot showing the flow of vOTU counts from viral families (left) to predicted host families (right) within the virulent fraction. Only vOTUs with both a viral family assignment and an iPHoP-derived host prediction are included; low-frequency families are grouped into ‘Others’. Ribbon width is proportional to the number of vOTUs linking each phage–host pair. **B)** Bipartite network of significant associations (Fisher’s exact test, *p* < 0.05) between viral families and predicted host families within the virulent fraction. Edge width is proportional to the Haldane–Anscombe-corrected odds ratio (OR). **C)** Alluvial plot as in A for the temperate fraction. **D)** Bipartite network as in B for the temperate fraction. Analyses were performed on the integrated community dataset (*n* = 4,357 vOTUs) across 30 pen-level samples.

Collectively, these results indicate that the swine nasopharyngeal phageome is organized into a small set of recurrent, host-linked taxonomic modules, with phage lifestyle systematically rebalancing the prominence of these modules across the community.

## Discussion

This study provides, to the best of our knowledge, the first field-scale, paired evaluation of DNA microbiome shotgun metagenomics (DNA-m) versus virus-like particle–enriched metagenomics (VLP-e) for swine nasopharyngeal specimens, and the first phageome catalogue derived from a respiratory site in any livestock species. By integrating both workflows across 30 paired samples from ten commercial farms (n = 60 libraries), we generate protocol-aware infrastructure and an ecological framework in which predicted replication strategy, host affiliation, and viral lineage converge as jointly constrained organizing axes of community structure.

Our paired benchmarking reveals that workflow selection is a key determinant of viral recovery in this niche. Since both workflows were applied to the same field-derived material, the paired design minimizes confounding by between-sample heterogeneity, so observed differences primarily reflect workflow-specific recovery [18,19]. The two approaches displayed a consistent trade-off: DNA-m favored assembly contiguity, while VLP-e maximized viral detection. This pattern is consistent with the design rationale of each approach: particle enrichment prioritizes signal-to-noise for viral discovery [17], whereas total DNA better supports assembly continuity and can capture cell-associated viral forms, including integrated prophages [19]. Critically, this complementarity extended beyond quantitative yield to community composition: overlap between workflows was minimal (<1%), and extraction method explained nearly half of the compositional variation (R² = 0.448), a partitioning that persisted when stratified by lifestyle. Comparable method-driven divergence has also been reported in soil virome–metagenome comparisons [17] and across multiple ecosystems [19], confirming that workflow choice materially alters ecological interpretation. However, in low-biomass settings, overlap estimates are inherently sensitive to detection thresholds and background-control stringency, so absence in one workflow should not be equated with biological absence but rather with workflow-dependent detectability near the limits of recovery. Cross-study comparisons in porcine respiratory viromics will therefore require transparent protocol reporting and, ideally, protocol-aware normalization.

By integrating both workflows under MIUViG quality tiers [20] and species-level dereplication, we constructed a curated, non-redundant catalogue of 2,501 representative vOTUs. The ≥50% completeness threshold prioritizes robustness over exhaustiveness, so the true viral diversity of this niche is likely considerably larger. Even under this conservative design, the catalogue exposes a striking degree of reference scarcity: most vOTUs lacked detectable analogues both at the nucleotide level against curated phage databases (94.8%) and at the protein-content level in the vConTACT2 network (59.2%), and a sizeable fraction of 146 VCs co-clustered exclusively with other catalogue members in lineages absent from public collections. Convergence of these methodologically independent metrics indicates that the recovered novelty reflects coherent lineages internal to the niche rather than fragmentary contigs. This gap is anatomically specific: dedicated catalogues are mature for the human and animal gastrointestinal tract [13,14,16,21], whereas respiratory virome studies in livestock have focused almost exclusively on eukaryotic pathogen detection [22–24]. Consistent with this reference-poor status, vOTU prevalence was strongly long-tailed and a sample-based accumulation curve did not approach saturation, indicating that nasopharyngeal phage diversity remains far from exhausted. The catalogue is therefore best framed as foundational, expandable infrastructure, with broader cohorts required to resolve rare lineages and test cross-population generalizability.

Within this catalogue, taxonomic resolution decreased sharply below the *Caudoviricetes* class, reflecting the well-documented lag between metagenomic phage discovery and formal classification [21,25,26]. To resolve lineage structure independently of database-anchored taxonomy, we combined two complementary scaffolds: the conserved single-marker terL, effective for delineating broad phage lineages [27,28], and vConTACT2 protein-content clustering [29], which approximates finer relationships. Both scaffolds were strongly nested, with vConTACT2 clusters consistently embedded within broader terL lineages (W_VC→terL_ = 0.86) and converged on a common architecture: a small number of dominant lineages flanked by a long tail of small or rare groups. This combination of multi-scale coherence and uneven lineage occupancy mirrors patterns reported for other host-associated phageomes [21,30] and provides a phylogenetically coherent, classification-independent scaffold for examining how replication strategy and host affiliation organize the nasopharyngeal phageome.

Using the sample-resolved community dataset (*n* = 4,357), we identified replication strategy as a major organizing axis of the swine nasopharyngeal phageome. Despite limited taxonomic resolution at lower ranks, predicted lifestyle explained a substantial proportion of compositional variation at both the family (R² = 0.406) and genus (R² = 0.612) levels. This supports the broader view that phageome structure stratifies along virulent–temperate fractions [11,12], and the signal was biologically coherent within the classified fraction: *Peduoviridae* were markedly temperate-enriched, consistent with the well-characterized temperate biology of P2-like phages [31], whereas virulent enrichment mapped to families dominated by lytic clades, including *Rountreeviridae*, *Schitoviridae*, and φ29-like *Salasmaviridae* [32–35].

This lifestyle-dependent structuring extended to predicted host range, generating a marked polarization across dominant upper-airway colonizers. Virulent phages were preferentially linked to *Bacteroidota*, particularly *Prevotella*, whereas temperate phages were predominantly associated with *Streptococcaceae* and *Moraxellaceae*. This polarization finds external support: population-scale gut virome surveys have shown that phages predicted to infect *Bacteroidota* hosts are predominantly virulent [36], while both *Streptococcaceae* and *Moraxellaceae* harbor abundant and structured prophage complements [37,38], consistent with the temperate enrichment observed here. Host assignability, however, was markedly asymmetric: temperate phages were substantially more likely to receive a host prediction than virulent phages (68.9% vs. 38.9%; OR = 3.49). This asymmetry likely reflects methodological rather than biological differences, since host-prediction tools rely on genomic signatures, such as compositional similarity and prophage-derived homology, that are inherently stronger for phages with a history of chromosomal integration [39], and the lifestyle-stratified host patterns should be interpreted within this context. Reassuringly, within-study CRISPR spacer–protospacer matches provided orthogonal, lifestyle-unbiased support for computational host predictions, with high concordance at the phylum (98.3%) and family (90.6%) levels in the supported subset.

The integration of viral taxonomy, host range, and lifestyle resolved a modular architecture in which a small number of recurrent, near-exclusive phage–host couplings concentrated the strongest associations: *Suoliviridae* linked to *Bacteroidota*, *Peduoviridae* to *Pseudomonadota*, and *Rountreeviridae* to *Bacillota*, consistent with the known tropism of these lineages in other ecosystems [31,32,40,41]. To the best of our knowledge, this is the first description of their convergence into a host-linked modular backbone within an upper respiratory phageome, with these couplings strengthening at finer host taxonomic resolution. Critically, these modules were conserved across lifestyles but differentially weighted by replication strategy: virulent communities emphasized the *Suoliviridae*–*Bacteroidota* axis and *Schitoviridae* links to *Moraxellaceae/Pasteurellaceae*, whereas temperate communities showed tighter coupling within *Peduoviridae*–*Pasteurellaceae* and *Aliceevansviridae*–*Streptococcaceae*. This differential weighting indicates that viral lineage, host affiliation, and replication strategy do not structure the nasopharyngeal phageome independently but act as jointly constrained ecological axes.

The ecological framework described above acquires particular relevance in the context of the post-weaning stage, a period during which the upper-airway bacterial community undergoes substantial reorganization [42,43]. Notably, *Streptococcaceae*, *Moraxellaceae*, and *Pasteurellaceae*, the three host families most prominently embedded within the modular phage–host architecture described here, are among the most affected taxa during this transition and harbor key opportunistic pathogens of the porcine respiratory disease complex (PRDC), including *Streptococcus suis*, *Glaesserella parasuis*, and *Pasteurella multocida* [2]. This convergence raises testable hypotheses: do bacterial shifts at weaning flow through linked phage lineages and alter the local lysis–lysogeny balance? Does this modulation, in turn, affect pathogen fitness during a critical epidemiological window? Resolving whether phages act as passive followers of bacterial dynamics or as active ecological modulators in this niche will require longitudinal and, ideally, interventional study designs.

In addition, from a translational standpoint, phage therapy against swine respiratory pathogens remains in its infancy, with only preliminary reports targeting *P. multocida* [44], *S. suis* [45], and *B. bronchiseptica* [46]. The catalogue and host–lineage associations generated here provide the first site-specific reference framework for this niche, enabling rational pre-selection of lytic candidates against respiratory colonizers. As niche-specific reference resources expand, reanalysis of this catalogue should progressively resolve annotation gaps and test the robustness of the modular architecture described here.

Several constraints should be considered when interpreting these patterns. First, nasopharyngeal swabs are intrinsically low-biomass specimens in which contaminating DNA can critically impact sequence-based analyses [47]; our workflow-matched blanks and conservative filtering reduce false positives but may also remove true low-abundance viruses, slightly reducing sensitivity near the detection limit. Second, pen-level pooling increases farm-level representativeness but masks inter-individual variation, so the patterns reported here reflect community-level averages rather than individual-animal dynamics. Third, short-read sequencing constrains genome closure and can fragment closely related viral variants, particularly those with high microdiversity or repetitive regions [48]; long-read or hybrid strategies were not feasible here because such libraries require substantially more input DNA than was recoverable from VLP-e preparations. Fourth, all ecological patterns derive from computationally predicted host and lifestyle annotations on a fraction of the recovered diversity; the uncharacterized majority may harbor signals that reinforce, attenuate, or restructure the patterns observed in the classified fraction [39,49]. Finally, increasing PCR cycles for low-input VLP-e libraries may bias coverage and relative representation [18], so quantitative contrasts should be interpreted cautiously even when inference is contig-based.

## Conclusions

Using a field-scale paired design, we showed that total-DNA and VLP-enriched metagenomics yield complementary, largely non-overlapping phage fractions from the swine nasopharyngeal niche. By integrating both workflows, we generate the first phageome catalogue and a marker-supported phylogenetic scaffold for a respiratory site in livestock. Predicted replication strategy emerged as a dominant organizing axis, with coherent lineage-level and host-affiliated shifts that converge into a reproducible modular architecture jointly constrained by viral lineage, host affiliation, and lifestyle. This catalogue and its associated ecological framework provide protocol-aware infrastructure and a scalable interpretable starting point for longitudinal, functional, and translational studies of the post-weaning porcine respiratory phageome.

## Methods

### Study design and farm sampling

Nasopharyngeal swab samples were collected in July 2025 from post-weaning piglets in ten commercial swine farms located in northwestern Spain (Castilla y León region). Farms operated under intensive production systems with conventional management practices. Within each farm, three pens were sampled. In each pen, five piglets aged 6-8 weeks were swabbed using deep nasopharyngeal insertion, and individual swabs were subsequently pooled at the pen level to generate one composite sample per pen. Overall, 150 piglets were sampled (*n* = 15 per farm), yielding 30 pooled pen-level samples (*n* = 3 per farm). Sampling was conducted under approval of the Ethics Committee of the University of León (ETICA-ULE-081-2025).

### Nasopharyngeal swab collection and dual-buffer design

Nasopharyngeal sampling was performed using aluminum-shaft viscose swabs (Deltalab, Spain). To enable direct comparison of extraction strategies while minimizing anatomical and procedural bias, sampling followed a symmetric crossover design using two preservation buffers: (i) 1 mL PBS 1× for total microbiota DNA extraction (hereafter DNA-m workflow), and (ii) 1 mL SM buffer (50 mM Tris-HCl pH 7.5, 100 mM NaCl, 8 mM MgSO₄) for virus-like particle enrichment (hereafter VLP-e workflow).

For each animal, two swabs were used to represent both nostrils and balance buffer order. One swab was used to sample the left nostril and then sequentially eluted into the DNA-m microtube (Deltalab, Spain) followed by the VLP-e microtube; a second swab was used to sample the right nostril with the inverse order (VLP-e first, then DNA-m). This procedure ensured that both nostrils contributed to both protocols while controlling for potential order effects.

Samples were transported refrigerated (4°C) to the BACRESPI laboratory (Animal Health Department, University of León, Spain) within 4h of collection, where they were pooled by pen and workflow to generate one DNA-m pool and one VLP-e pool per pen. The final dataset comprised 60 pooled samples: 30 DNA-m and 30 VLP-e, corresponding to the 30 pen-level sampling points.

### Nucleic acid extraction workflows

#### Total DNA extraction (DNA-m workflow)

Pen level DNA-m pools were processed using the QIAamp DNA Microbiome Kit (QIAGEN, Germany) following the manufacturer’s protocol. Briefly, this workflow combines selective lysis and depletion of host (eukaryotic) DNA, followed by mechanical disruption of microbial cells by bead beating and in silica column purification. DNA-m was eluted in a final volume of 60 µL and stored at −80 °C until use.

#### Virus-like particle enrichment and viral DNA extraction (VLP-e workflow)

VLP enrichment and viral DNA extraction were adapted from Shkoporov et al. [50] with slight modifications for low-biomass respiratory swab pools. Briefly, SM-buffer VLP-e pen-level pools were clarified by two sequential low-speed centrifugations (3,200 g for 10 min at 4°C), and the resulting supernatant was passed twice through 0.45-µm cellulose acetate syringe filters (Labbox, Spain) to deplete residual cells and large cellular debris while retaining virus-sized particles.

Viral particles were concentrated by polyethylene glycol (PEG) precipitation: NaCl (Panreac, Spain) and PEG-8000 (Fisher Scientific, NJ, United States) were added to the clarified supernatant to reach final concentrations of 0.5 M NaCl and 10% (w/v) PEG-8000, and the mixture was incubated overnight (∼16 h) at 4 °C. Pellets were collected by centrifugation (3,200 g for 20 min at 4°C), resuspended in 400 µL SM buffer, and subjected to chloroform extraction to remove residual lipids and membrane-associated contaminants. An equal volume of chloroform (Sigma-Aldrich, MO, United States) was added, the mixture was vortexed vigorously for 30 s, and phases were separated by centrifugation (2,500 × g, 5 min, room temperature). The aqueous phase was carefully recovered and subjected to an additional brief centrifugation and transfer step to minimize chloroform carryover.

To remove free (non-encapsidated) nucleic acids, samples were supplemented with 40 µL of a solution containing 10 mM CaCl₂ and 50 mM MgCl₂ to support nuclease activity. Free DNA was digested with TURBO DNase (8 U; Invitrogen, Thermo Fisher Scientific, Lithuania) and free RNA with RNase I (20 U; Thermo Fisher Scientific, Lithuania) in the presence of 1× TURBO DNase buffer. The mixture was incubated with gentle agitation for 1 h at 37 °C, followed by heat inactivation at 70 °C for 10 min.

Protein digestion and capsid lysis were then performed by adding Proteinase K (40 µg; QIAGEN, Germany) and 20 µL of 10% sodium dodecyl sulfate (SDS; Sigma-Aldrich, MO, United States), followed by incubation with 100 µL of phage lysis buffer (4.5 M guanidinium isothiocyanate, 44 mM sodium citrate pH 7.0, 0.88% sarkosyl, 0.72% 2-mercaptoethanol) at 65°C for 10 min. Viral DNA was purified by two sequential phenol–chloroform–isoamyl alcohol (25:24:1, pH 8; Fisher Scientific, NJ, United States) extractions, followed by centrifugation at 8,000×g for 5 min at room temperature. The final aqueous phase was cleaned using the DNeasy Blood & Tissue Kit (QIAGEN, Germany) according to the manufacturer’s protocol, with a final elution volume of 60 µL. DNA was stored at −80 °C until use.

Given the expected low DNA yields from VLP-enriched respiratory specimens, one negative extraction control per workflow (DNA-m and VLP-e) was processed alongside each extraction batch using the same reagents and procedures but without sample input, allowing systematic monitoring of potential reagent-derived or environmental contamination throughout the extraction and library preparation processes.

### Library preparation and short-read sequencing

Extracted DNA from both workflows was quantified using the Qubit dsDNA High Sensitivity assay (Invitrogen, Thermo Fisher Scientific, MA, USA). Sequencing libraries were prepared using the TruSeq DNA Nano kit (Illumina, CA, USA) following the manufacturer’s recommendations, with a target insert size of approximately 350 bp. For VLP-e extracts with DNA amounts below recommended input, PCR amplification cycles during library preparation were increased from 8 to 16 to obtain sufficient library yield. This modification was applied to 18 of the 30 VLP-e libraries. Although increasing PCR cycles can introduce amplification bias affecting coverage evenness and relative representation [18], this step was necessary to generate sequenceable libraries from the low DNA quantities recovered after VLP enrichment of respiratory specimens. Downstream ecological inference in this study relies primarily on assembled contigs and dereplicated vOTUs rather than on read-level quantification, which partially mitigates amplification-driven biases.

Fragment size distributions were assessed using a 5200 Fragment Analyzer System (Agilent Technologies, CA, USA), and final library concentrations were measured by Qubit dsDNA High Sensitivity assay prior to sequencing. Paired-end sequencing (2 × 150 bp) was performed on an Illumina NextSeq 500 platform (Illumina, CA, USA) using a NextSeq 500/550 High Output kit v2.5 (300 cycles) (Illumina, CA, USA), following standard Illumina protocols. Libraries were multiplexed to target approximately 6.6 Gbp of raw data per sample. However, several VLP-e libraries yielded lower sequencing depth than initially targeted. Prior to processing the full sample set, both workflows were applied to a pilot subset of samples to confirm adequate DNA yield, library performance, and sequencing quality.

### Bioinformatic processing

#### Read preprocessing, quality control and viral enrichment metrics

Raw paired-end reads were processed using fastp (v.1.0.0) [51] for adapter trimming, quality assessment and quality filtering using default parameters. Summary quality metrics were inspected across runs to ensure consistent read quality between extraction workflows. Quality-filtered reads were additionally assessed with ViromeQC (v1.0.2) [52] with default settings to estimate viral enrichment scores and residual contamination from ribosomal RNA (SSU and LSU) and single-copy bacterial marker genes. Per-sample reports were collated to compare enrichment metrics between workflows.

Computational removal of host-derived reads was not performed prior to assembly. The QIAamp DNA Microbiome Kit (DNA-m) and the filtration–nuclease treatment (VLP-e) both substantially deplete eukaryotic DNA during extraction. Because all downstream analyses operate at the contig level, any residual host-derived reads that co-assemble into non-viral contigs are effectively excluded during post-assembly viral classification.

#### Metagenome assembly and phage genome reconstruction

Quality-filtered reads were assembled de novo on a per-library basis using MEGAHIT (v1.2.9) [53] with the default iterative k-mer strategy (k = 21, 29, 39, 59, 79, 99, 119, 141) and a minimum contig length threshold of 500 bp. This length threshold was chosen to be consistent with the expected genome architecture of bacteriophages and to reduce the retention of spurious short fragments.

Phage genome reconstruction was subsequently performed per sample using Phables (v1.4.1) [54]. Briefly, MEGAHIT graph outputs were converted to GFA format using the megahit_toolkit contig2fastg utility, and quality-filtered paired reads were mapped back to the assembly graph to provide coverage information. Phables was then run per sample to resolve phage genome paths from fragmented assembly components, producing both resolved phage genomes and unresolved viral edges. Per-sequence coverage estimates for all reconstructed sequences were computed using koverage v0.0.19 [55].

#### Decontamination using negative extraction controls

Given the low DNA input inherent to nasopharyngeal microbiome, particularly in VLP-e libraries, and the consequent susceptibility to reagent-derived and background contamination [47], Phables-derived contigs were decontaminated using batch- and workflow-matched extraction-negative controls. Specifically, negative-control reads were mapped against the corresponding sample assemblies using KMA (v1.4.0) [56]. Contigs with any detectable read mapping from negative controls were conservatively removed prior to all downstream analyses.

#### Viral sequence identification and quality assessment

Putative viral sequences were identified from the decontaminated Phables outputs using two complementary classifiers run independently. VirSorter2 (v2.2.4) [57] was run with the virome-specific workflow, including dsDNAphage and ssDNA groups and a minimum score threshold of 0.5. In parallel, geNomad (v1.11.0) [58] was run in end-to-end mode. Viral calls from each tool were independently evaluated using CheckV (v1.0.3) [59] in end-to-end mode to estimate completeness and host contamination, and to assign standard quality categories (*i.e.*, complete, high-quality, medium-quality, low-quality, or not-determined). Outputs from both classifiers were retained for downstream integration and cross-tool concordance checks.

#### Non-redundant nasopharyngeal phage catalogue construction

Viral predictions obtained from geNomad and VirSorter2 were integrated at the per-sample level. For contigs independently identified by both classifiers, redundant records were resolved by retaining the prediction with the lowest CheckV-estimated contamination, yielding a non-redundant set of viral contigs per library. Quality filtering was then applied using CheckV completeness estimates, retaining only sequences with ≥50% estimated completeness (i.e., medium-quality or higher), according to MIUViG standards [20].

All retained viral sequences were pooled across samples and dereplicated into viral operational taxonomic units (vOTUs) using a two-stage clustering strategy. In the first stage, candidate clusters were generated by pre-clustering with Mash (v2.3) [60] using a k-mer size 21 and a sketch size 10,000; sequence pairs with a Mash distance ≤0.05 were linked into candidate clusters. In the second stage, clusters were refined by pairwise nucleotide alignment with MUMmer4 v4.0.1 [61] using nucmer (--maxmatch) followed by delta-filter (-q -r), and sequences sharing ≥95% average nucleotide identity (ANI) were assigned to the same vOTU, consistent with the species-level demarcation threshold widely adopted in viral metagenomics [20]. Within each vOTU, the representative sequence was selected by prioritizing highest CheckV-estimated completeness, then lowest contamination, and finally longest contig length. The resulting set of representative sequences constitutes the non-redundant swine nasopharyngeal phage catalogue released with this study.

To assess phylogenetic coherence within the curated vOTU catalogue, the large terminase subunit (terL) was identified and extracted with Pharokka v1.7.5 [62]. Only terL amino-acid sequences of ≥100 residues were retained for phylogenetic analysis. Retained terL sequences were aligned with MAFFT v7.471 [63] with the following options: --auto --maxiterate 1000 --retree 2 --adjustdirection. The resulting alignment was trimmed with TrimAl v2.0 [64] using a gap-threshold of 0.1 to remove poorly aligned positions. A maximum-likelihood phylogeny was inferred with IQ-TREE2 v2.4.0 [65] under the LG+G4 substitution model, with branch support assessed by 1,000 ultrafast bootstrap replicates and 1,000 SH-aLRT replicates. Proteins identified from Pharokka were input to analysis with vConTACT2 v.0.11.3 [29] using the provided ProkaryoticViralRefSeq211-Merged reference database with default parameters (--rel-mode Diamond). Taxonomic assignments for reference genomes were retrieved from the database metadata as provided by vConTACT2. The resulting clustering from vConTACT2 and terL were compared using http://www.comparingpartitions.info/ [66] providing adjusted Wallace coefficient for direction-dependent agreement and the adjusted Rand index (ARI)for global congruence.

Representative vOTUs were screened against the PhageScope sequence collection [67] using MMseqs2 v17-b804f [68] in nucleotide search mode. MMseqs2 outputs were parsed to retain alignment-level similarity metrics, including fraction of identical nucleotides (fident), query coverage (qcov; fraction of the vOTU sequence covered by the alignment), and target coverage (tcov; fraction of the matched PhageScope sequence covered), together with e-values. Hits were first subjected to a permissive pre-filter (e-value ≤ 1e−5, fident ≥ 0.50, qcov ≥ 0.20) to remove weak or short matches, after which the best hit per query was refined by nucleotide alignment with MUMmer4 (nucmer --maxmatch, delta-filter -1, show-coords) and selected prioritizing highest percent identity and alignment length. Similarity to previously described phages was reported at two stringency levels. A permissive match required fident ≥ 0.90, qcov ≥ 0.25, tcov ≥ 0.20, aligned length ≥ 2 kb, and matched target length ≥ 2 kb. A stringent (high-confidence) match additionally required fident ≥ 0.92, qcov ≥ 0.35, aligned length ≥ 5 kb, target length ≥ 5 kb, and either tcov ≥ 0.50 or a query length ≥ 15 kb. Unless otherwise stated, “close matches” refer to the lenient criterion, with the stringent subset reported separately.

#### Taxonomy, lifestyle and host assignment of vOTUs

Taxonomic classification and replication strategy (lifestyle) inference were performed on the quality-filtered, sample-resolved community dataset (i.e., all vOTUs with CheckV-estimated completeness ≥50% detected across both extraction protocols) to preserve ecological variation across samples. Viral taxonomy and lifestyle were assigned using PhaBOX2 (v2.1.11) [69] in end-to-end mode with a minimum contig length of ≥500 bp, using the summary outputs. Lifestyle predictions were derived from the integrated phaTYP module, and taxonomy fields reported by PhaBOX2 were used for downstream summaries and enrichment analyses. Summary tables integrating vOTU identifiers, CheckV-derived quality metrics, taxonomic assignments and lifestyle classifications were generated for all downstream ecological analyses.

Putative bacterial hosts were predicted from vOTU sequences using iPHoP (v1.3.3) [39]. When multiple host predictions were returned for the same vOTU, the prediction with the highest iPHoP confidence score was retained as the single host assignment and integrated into the summary tables.

#### Independent validation of host predictions by CRISPR spacer analysis

To provide orthogonal, within-study support for computational host predictions, CRISPR spacer–protospacer matching was performed. Bacterial contigs ≥1,000 bp were extracted from the decontaminated assemblies and taxonomically classified using Kraken2 v2.1.3 [70] against the Genome Taxonomy Database (GTDB) v220.0 [71].

CRISPR arrays were detected on the taxonomically assigned bacterial contigs using CRISPRCasFinder v2.0.3 [72], with a minimum evidence level of 2 and a minimum of two spacers per array. Spacer sequences of 25–60 nt were extracted and aligned against the viral sequence set using BLASTn v2.16.0+ [73] with the blastn-short task, a word size of 7, a minimum percent identity of 90%, an e-value threshold of 1 × 10⁻³, and retaining only the best hit per spacer. Resulting alignments were subsequently filtered to retain only spacer-like matches meeting stringent criteria: percent identity ≥95%, ≤1 mismatch, no gaps, alignment length between 25 and 60 nt, and target viral contig with CheckV-estimated completeness ≥50%. Spacer–protospacer matches passing these thresholds were used to independently assess concordance with iPHoP-derived host predictions and to evaluate whether the lifestyle distribution of CRISPR-supported phages differed from that of the full dataset.

### Statistical analyses and figure visualization

All downstream analyses reflected the paired sampling design and the compositional nature of virome count data. Statistical tests were two-sided and performed in R v4.4.3 [74]. Multiple testing was controlled using the Benjamini–Hochberg false discovery rate (FDR) procedure [75], with significance defined at adjusted *p* < 0.05. Figures were produced with ggplot2 v4.0.1 [76] and related extensions, and finalized in Inkscape v1.4.2 (https://inkscape.org/).

#### Protocol-dependent recovery metrics

Protocol-dependent differences in sequencing yield, ViromeQC enrichment scores, residual rRNA and bacterial marker gene contamination, assembly statistics, and viral-contig recovery were compared across matched pen-level pairs using paired Wilcoxon signed-rank tests. Associations between ViromeQC viral enrichment scores and the number of recovered viral contigs were assessed using Spearman’s rank correlation.

#### Compositional analysis of community structure

Extraction-method effects on viral community structure were assessed using a compositional data analysis framework. vOTU abundance matrices were prepared by adding a pseudocount of 0.5 to all entries and applying the centered log-ratio (CLR) transformation. Aitchison distances were computed as Euclidean distances in CLR-transformed space. Differences in community composition were tested by permutational multivariate analysis of variance (PERMANOVA) using the adonis2 function in the R package vegan v2.7.2 [77]. Homogeneity of multivariate dispersions was evaluated using vegan::betadisper. Community ordination was visualized by principal coordinates analysis (PCoA) on Aitchison distances, and all compositional analyses were repeated after stratification by predicted phage lifestyle (virulent vs temperate).

#### Phylogenetic characterization of the curated database

Phylogenetic characterization of the curated database used the terL marker. The IQ-TREE2 maximum-likelihood terL tree was imported into R and visualized as a circular phylogeny with concentric annotation rings using ggtree v3.14.0 [78]. Discrete lineages were defined by hierarchical clustering of cophenetic distances using average-linkage agglomeration and the cutree function with k = 19. Enrichment of taxonomic, lifestyle, and predicted host annotations within each terL cluster was tested using two-sided Fisher’s exact tests. Odds ratios (OR) were computed using the Haldane–Anscombe (HA) correction (adding 0.5 to all cells of the contingency table), and 95% confidence intervals (95% CI) were derived from the corrected tables.

Sample-based vOTU accumulation curves were generated by random permutation of sample order, and observed cumulative richness was fitted to two alternative population models: a power-law function (no upper asymptote, consistent with an open viral gene pool) and an asymptotic exponential function (consistent with a closed, fully sampled diversity). Model selection was performed by AIC.

#### Ecological compositional analyses

Ecological compositional analyses of the sample-resolved community dataset were performed on vOTU abundance matrices aggregated at successive taxonomic levels: viral taxonomy (class, family, genus) and predicted host taxonomy (phylum, family, genus), stratified by predicted lifestyle. Using the same CLR/Aitchison framework described above, lifestyle-associated shifts in viral taxonomic and host-affiliation profiles were tested by PERMANOVA and visualized by PCoA.

Viral and host taxa differentially associated with virulent or temperate fractions were identified using paired Wilcoxon signed-rank tests on CLR-transformed abundances, with effect sizes summarized as median paired differences (Δ). Differences in host-assignment rates between lifestyles were assessed using Fisher’s exact tests. Host–phage taxonomic coupling (viral family/genus vs. host phylum/family/genus) was quantified using Fisher’s exact tests, reporting HA-corrected OR with 95% CI. Association patterns were visualized using alluvial plots to summarize dominant phage–host flows, heatmaps of effect sizes across taxa, and bipartite networks highlighting significant links.

## Declarations

### Ethics approval and consent to participate

All procedures involving animals were approved by the Ethics Committee of the University of León (Reference Number ETICA-ULE-081-2025).

### Consent for publication

Not applicable.

### Availability of data and materials

The DNA sequences from the 60 metagenomic libraries from 10 Spanish swine farms and the ten DNA extraction controls, are available in the NCBI Sequence Read Archive (SRA) repository under accession number PRJNA1467997. The catalogue containing 2,501 vOTUs is also available in https://doi.org/10.5281/zenodo.20119467.

### Authors’ contributions

Study design was performed by OMA, SMM, CBGM, BM, CD, and JG. Experiment was conducted by OMA, and JG, with support of SMM and CD. Laboratory analyses were performed by JG, SMM and OMA. Bioinformatic and statistical analyses were performed by OMA, JG and CD. SMM, CD, and BM provided technical and scientific support for the analysis. Funding was provided by BM, SMM and, OMA. OMA, CD and JG participated in the manuscript writing or contributed to its revision. All authors revised the manuscript and approved the final version.

## Acknowledgments

We acknowledge the excellent technical assistance provided by Ana Isabel Pastor, Lucía Manzanares Vigo and Mario de Frutos in sampling collection, as well as the help provided by Katharina Thomas in the sequencing library preparation. We would like to thank also the veterinary practitioners and farmers for their willingness to collaborate.

## Additional files

**Additional file 1: Tables S1-S3.** Summary and statistical analysis of sequencing, quality-control, assembly and viral enrichment metrics for DNA microbiome (DNA-m) and virus-like particle–enriched (VLP-e) extraction workflows. Paired Wilcoxon signed-rank test statistics and Benjamini–Hochberg-adjusted *p*-values are provided for all metrics. *n* = 60 libraries (30 DNA-m + 30 VLP-e). File format: .xlsx

**Additional file 2: Figure S1. Construction of a curated, non-redundant swine nasopharyngeal phage catalogue. A)** UpSet plot of viral contigs independently identified by geNomad and VirSorter2, coloured by CheckV quality tier. The vertical bars show intersection sizes; the horizontal bars (left) show total set sizes per classifier; the dot matrix below indicates intersection membership. **B)** Number of representative vOTUs detected per pen-level sample. **C)** Sample-based vOTU accumulation curve. Black dots show observed cumulative vOTU richness; lines indicate the fitted power-law (red) and asymptotic exponential (blue) models, extrapolated to *n* = 40 samples. The curated catalogue comprises *n* = 2,501 representative vOTUs across 30 pen-level samples.

**Additional file 3: Tables S4–S6.** Annotation and enrichment analysis (*p* < 0.05) of terL phylogenetic clusters (*k* = 19) within the curated vOTU catalogue (*n* = 2,501). File format: .xlsx

**Additional file 4: Tables S7–S9.** Protein-content–based clustering of the curated vOTU catalogue with reference phage genomes using vConTACT2, and concordance with terL phylogenetic clusters. Tables include the composition of each viral cluster (VC) in terms of catalogue vOTUs and reference genomes, the per-vOTU assignment to VC and terL cluster with concordance categorization, and the pairwise mapping between VCs and terL clusters. *n* = 2,501 representative vOTUs distributed across 392 VCs of ≥2 members. File format: .xlsx

**Additional file 5: Tables S10–S15.** Detailed summary of viral taxonomic composition and lifestyle-associated differential abundance at family and genus levels. Tables include the distribution of vOTUs across viral families and genera in the integrated community dataset, stratified by predicted lifestyle, and paired Wilcoxon signed-rank test results for lifestyle-associated shifts in CLR-transformed abundances, reporting median paired differences (Δ) and adjusted *p*-values. *n* = 4,357 vOTUs across 30 pen-level samples. File format: .xlsx

**Additional file 6: Figure S2. Viral and host taxonomic composition at the genus level. A)** Stacked bar plot of the proportion (%) of vOTU counts classified by viral genus per sample. **B)** Stacked bar plot of the proportion (%) of predicted host genera (iPHoP) of vOTU counts per sample, excluding vOTUs without host assignment. In both panels, samples are ordered by farm and pen, and low-frequency genera are grouped into ‘Others’. Analyses were performed on the integrated community dataset (*n* = 4,357 vOTUs) across 30 pen-level samples. File format: .pdf

**Additional file 7: Tables S16–S18.** Lifestyle-dependent host composition at phylum, family and genus levels. For each taxonomic rank, paired Wilcoxon signed-rank test results are reported for differences in CLR-transformed host abundances between virulent and temperate fractions, including the number of paired observations, median paired differences (Δ), and adjusted *p*-values. *n* = 4,357 vOTUs across 30 pen-level samples. File format: .xlsx

**Additional file 8: Tables S19-S22.** CRISPR spacer–protospacer analysis for independent validation of host predictions. *n* = 65 vOTUs supported by CRISPR spacer matches; 58 with both CRISPR and iPHoP assignments. File format: .xlsx

**Additional file 9: Tables S23–S25.** Enrichment analysis of phage–host taxonomic associations at the phylum level (viral family vs host phylum). File format: .xlsx

**Additional file 10: Tables S26–S28.** Enrichment analysis of phage–host taxonomic associations at the family level (viral family vs host family). File format: .xlsx

**Additional file 11: Tables S29–S31.** Enrichment analysis of phage–host taxonomic associations at the genus level (viral family vs host genus). File format: .xlsx

