## Supplementary figures and images for "Paired viromics resolves the modular ecological architecture of the swine nasopharyngeal phageome"

### Additional file 2

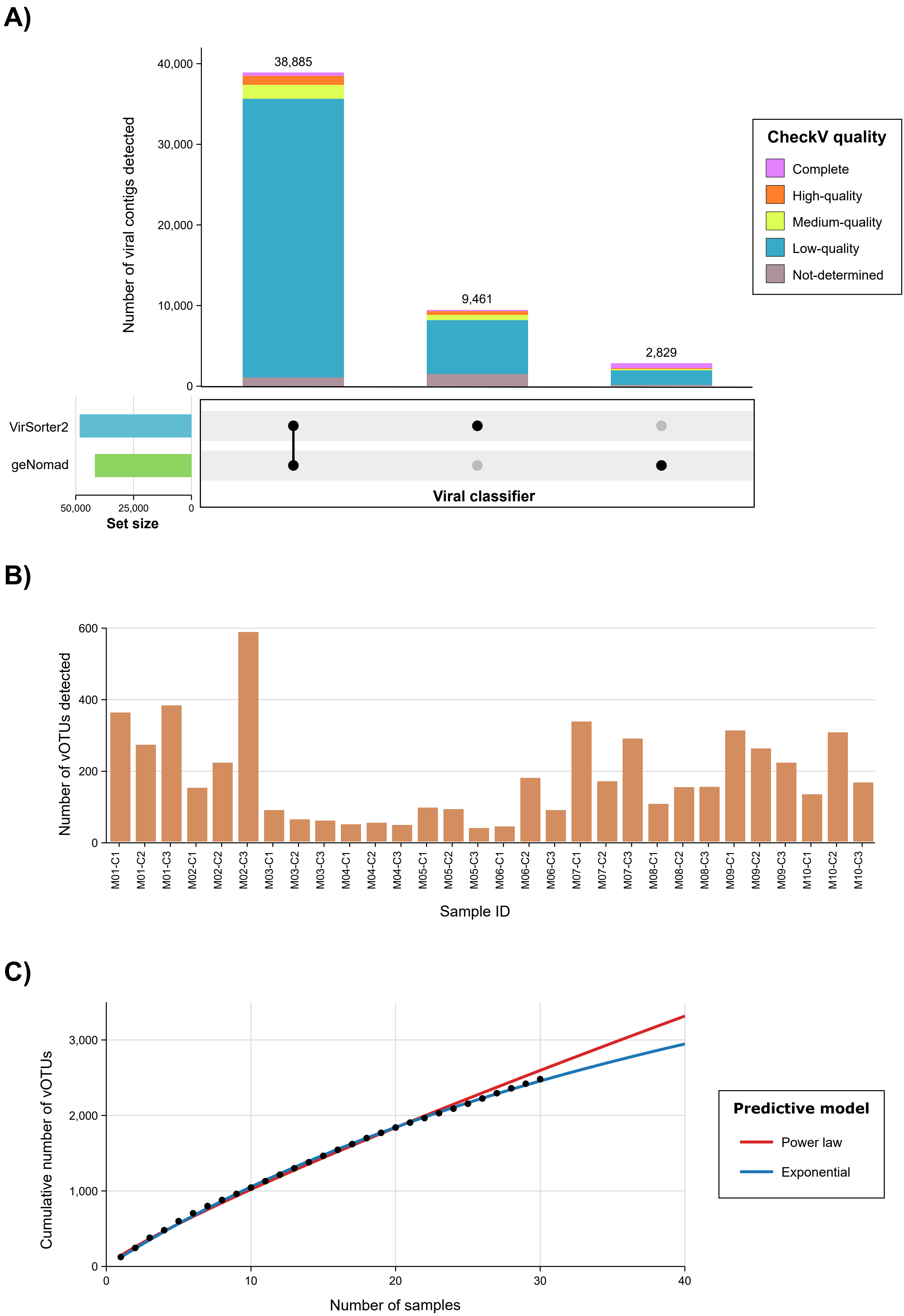

### Additional file 6

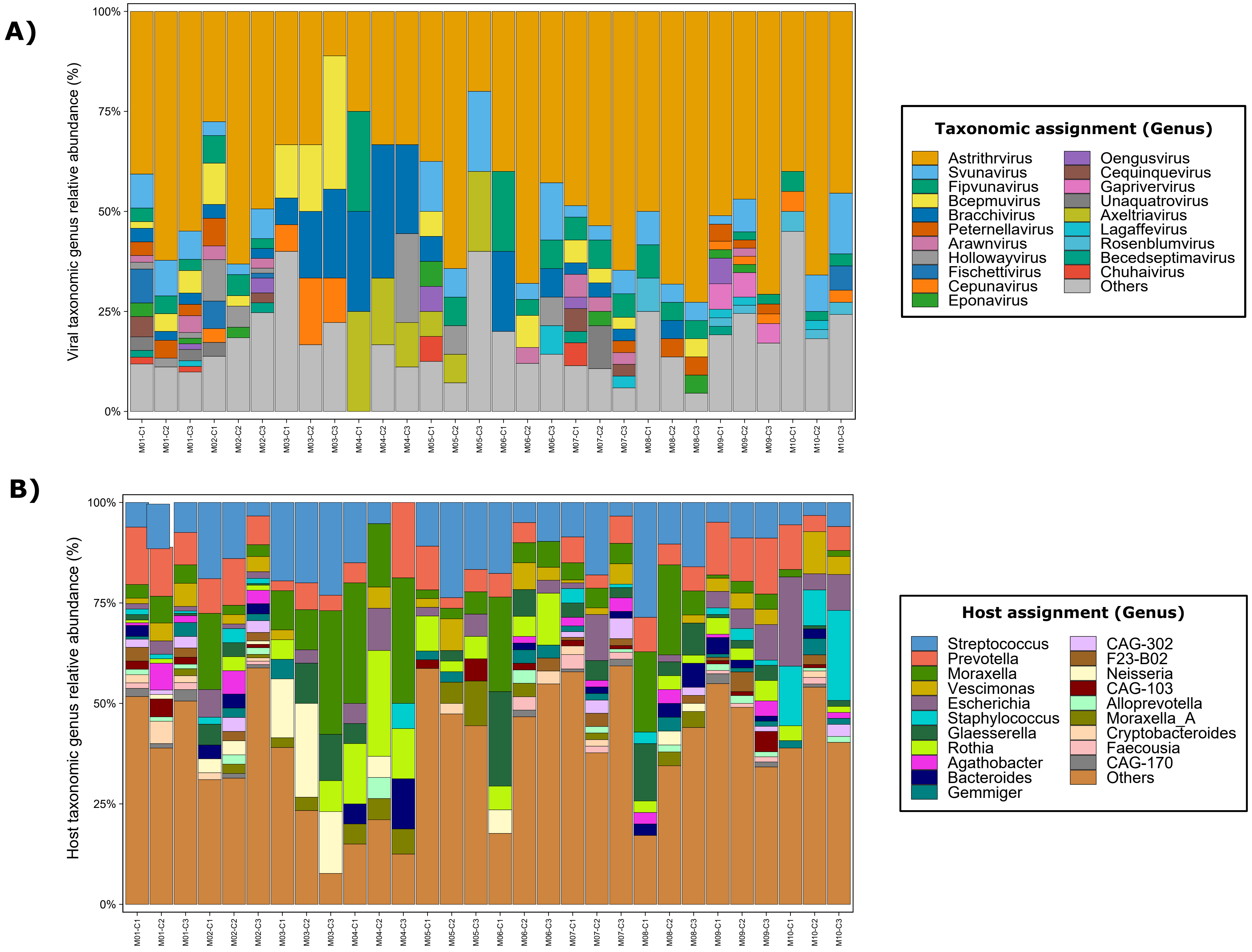
